# The Latentverse: An Open-Source Benchmarking Toolkit for Evaluating Latent Representations

**DOI:** 10.1101/2025.04.25.650676

**Authors:** Yoanna Turura, Samuel Freesun Friedman, Aurora Cremer, Mahnaz Maddah, Sana Tonekaboni

## Abstract

Self-supervised representation learning is a powerful approach for extracting meaningful features without relying on large amounts of labeled data, making it particularly valuable in fields like healthcare. This enables pretrained models to be shared and fine-tuned with minimal data for various downstream applications. However, evaluating the quality and behavior of these representations remains challenging. To address this, we introduce Latentverse, an open-source library and web-based platform for evaluating latent representations. Latentverse generates detailed reports with visualizations and metrics that provide a comprehensive perspective on different properties of representations, such as clustering, disentanglement, generalization, expressiveness, and robustness. It also allows for the comparison of different representations, enabling developers to refine model architectures and helping users assess how well an embedding model aligns with the requirements of their specific applications.

**Data and Code Availability:** The Latentverse code is available at: https://github.com/broadinstitute/ml4h-latentverse.

**Institutional Review Board (IRB):** This work doesn’t require IRB approval.

## 1. Introduction

Self-supervised representation learning (SSL) and more recently foundation models (Bommasani et al., 2021) have become a popular approach for extracting generalizable and informative features from data without relying on extensive manual annotation (Gui et al., 2024). Such approach is particularly beneficial in healthcare, where data annotation is both expensive and labor-intensive (Krishnan et al., 2022). These models have significantly outperformed task-specific architectures in settings with limited labeled data (Wornow et al., 2023), while also improving domain adaptation and robustness to distribution shifts (Guo et al., 2024). This has resulted in an increase in sharing pre-trained representation learning models that enable customization with minimal labeled data.

Despite this success, it still remains difficult for users to determine which pre-trained representations best suit their specific applications. Different methods incorporate distinct inductive biases and training strategies, leading to representations that capture varying aspects of the data (Bengio et al., 2013). Some models emphasize semantic consistency, while others prioritize feature disentanglement, robustness to distribution shifts, or domain adaptation (Locatello et al., 2019; Higgins et al., 2018). This variability makes it difficult to determine which pretrained model is best suited for a specific application, particularly in healthcare, where representations must retain clinically meaningful information. Beyond training models, embedding analysis can also serve as a powerful tool for knowledge discovery. By exploring embedding spaces, researchers can gain insights into dataset structure, identify patterns, or even potential challenges and biases in the embeddings. Developing a comprehensive evaluation framework is essential to help downstream users assess the suitability of pre-trained representations for their specific applications. Such a tool should systematically evaluate key properties, ensuring that learned representations align with the requirements of real-world tasks and would enable users to make informed decisions when selecting and fine-tuning those pre-trained models for their applications.

In this work, we introduce Latentverse, an opensource library and a web-based platform for comprehensive assessment of the quality of latent representations. Latentverse provides a suite of evaluation metrics covering clustering performance, disentanglement, and robustness to perturbations, offering insights into key properties such as expressiveness, separability, and stability. Users can use Latentverse to benchmark their models or compare different embedding strategies for their data. By uploading only learned embeddings—without requiring access to raw data—Latentverse ensures privacy-preserving evaluation with no concerns about sensitive information exposure. We also showcase the utility of Latentverse for evaluating the quality of clinical representations obtained from different medical modalities.

By sharing Latentverse as an open-source tool, we aim to foster community contributions, making it a widely adopted resource for evaluating latent representations across various domains.

## 2. Evaluating Latent Representations

Several tools have been developed for analyzing and interpreting learned representations, each with a different focus. In language models, interpretability has driven interest in latent space exploration. Unveiling LLMs (Bronzini et al.) examines how LLMs encode and process knowledge across layers using activation patching and dynamic knowledge graphs. While this provides insight into model reasoning, it is primarily designed for factual verification rather than structured representation evaluation.

Other tools focus on domain-specific representation analysis. Latent Space Explorer (Cecconello et al., 2022) applies unsupervised learning to identify patterns in large-scale astrophysical datasets, leveraging techniques like autoencoders and contrastive learning. However, it is designed for visualization rather than structured embedding evaluation. Similarly, Emblaze (Kwon et al., 2022) facilitates interactive embedding exploration through animated scatter plots and neighborhood analysis in computational notebooks, but it lacks a quantitative framework for assessing representation quality. Multimodal Latent Space Explorer (Kwon et al., 2023) specifically targets multimodal representation learning, helping researchers compare embeddings across modalities like MRI scans and ECG signals. Our objective is to provide a structured, general-purpose evaluation framework that quantitatively measures representation quality while offering an accessible web interface, enabling users to assess embeddings without requiring data sharing.

There are many ways to evaluate the quality of learned representations, and different metrics offer insights into various aspects. While there is no single definition of what makes a representation *good*, these metrics are invaluable for understanding their strengths and weaknesses.

We surveyed the literature to compile a comprehensive list of evaluation metrics that offer a holistic view of representation quality. This section outlines the various categories of properties that Latentverse evaluates and details the selection of metrics implemented to quantify different aspects of representation quality. We envision Latentverse as an evolving tool, constantly integrating novel metrics, and enabling users to extend and enhance its capabilities over time as new evaluation approaches emerge.

### 2.1 Downstream generalization

A key advantage of using latent representations of pretraining SSL models is their ability to transfer learned knowledge for training models on limited labeled data across various tasks Del Pup and Atzori (2023). Therefore, one of the most common evaluation of representations is measuring how well they support different downstream tasks, often referred to as probing (Belinkov, 2022). By training supervised models to predict labels from learned embeddings, probing tests offer insights into how well representations encode task-relevant information (Hewitt and Manning, 2019; Devlin et al., 2019). Since they require labeled data, these tests also help assess the generalization of representations across tasks.

When probing representations, there is a trade-off between model complexity and performance. Some advocate for using complex models to probe representations (Pimentel et al., 2020), while others emphasizes the importance of simpler models for better interpretability (Voita and Titov, 2020). As a result, the probing test in Latentverse is designed to explore this trade-off by measuring performance for probing models with varying complexity levels (Ferreira et al., 2021), helping users assess how well representations generalize across diverse tasks.

### 2.2 Clusterability

Another quality of representations can be measured by how well the data naturally forms distinct clusters, providing both quantitative and visual insights into the latent space structure. The clusterability test characterizes both the intrinsic geometric structure of the latent space and the neighborhood relationships between data points, offering insights into how well SSL models capture semantic similarities (Xie et al., 2019; Schneider et al., 2023). Meaningful clustering structures in the latent space often indicate representations that are well-aligned with the true underlying data distribution, leading to better downstream performance and generalization (Zhao et al., 2023; Caron et al., 2020). Several metrics have been proposed to quantify clusterability, each capturing different aspects of structure and separation. Latentverse integrates the following key clusterability metrics:

(1) The **Silhouette Score** (Rousseeuw, 1987) measures how similar a data point is to its own cluster *a*(*i*) compared to other clusters *b*(*i*), with values ranging from 1 (poor clustering) to *−*1 (perfect clustering). (2) The **Davies-Bouldin Index (DBI)** (Davies and Bouldin, 1979) quantifies the average similarity ratio between each cluster and its most similar cluster, with a lower value indicating better-defined clusters. (3) The **Normalized Mutual Information (NMI)** (Danon et al., 2005) score that measures the similarity between clustering results and ground truth labels. It ranges from 0 (no shared information) to 1 (perfect alignment). NMI quantifies how much information is retained between the predicted cluster assignments and the true labels, normalized by their entropy. This metric is widely used in clustering evaluation to ensure that clusters align well with known categories.

### 2.3. Disentanglement

Disentangling factors of variation in data is another key property of meaningful representations. Disentanglement metrics measure how well a representation separates underlying factors of variation in the data. Latentverse implements widely used metrics from two main categories:(1) Predictor-based metrics that use regressors or classifiers to predict factors from latent dimensions. While these metrics are effective for both continuous and categorical factors, they require careful model selection and tuning. (2) Information-based metrics estimate mutual information between latent codes and factors, making them more parameter-efficient and assumption-light.

The **DCI** (Eastwood et al.) metric evaluates the quality of latent representations based on three key measures: (1) Disentanglement: Assesses whether each generative factor is captured by a single latent dimension. A higher score means that individual factors are encoded in separate dimensions rather than being entangled across multiple ones. (2) Completeness: Evaluates how well each generative factor is captured by a small subset of latent dimensions. A high score indicates that information about a factor is not spread across many dimensions but is concentrated in a few. (3) Informativeness: Measures how well the learned representations predict the original generative factors. Together, these measures provide a comprehensive assessment of how structured and interpretable a representation is.

The Separated Attribute Predictability **(SAP)** (Kumar et al., 2018) score is a metric designed to evaluate the degree of disentanglement in a learned latent representation. It measures how well each latent variable can predict individual generative factors using simple linear regression (for continuous factors) or threshold-based classification (for categorical factors). It is computed by constructing a matrix of predictive scores and selecting the best-performing latent for each factor. A higher SAP score indicates better disentanglement, as each generative factor is more easily predictable from a single dimension.

The Mutual Information Gap **(MIG)** (Chen et al., 2018) is another metric designed to measure the degree of disentanglement in a learned latent representation by assessing the alignment between latent variables and a single generative factor. It quantifies how well the most informative latent dimension captures the factor while ensuring minimal redundancy across other dimensions. Specifically, MIG computes the difference in mutual information between the most and the second-most predictive latent dimension, normalized by the entropy of the generative factor. A higher MIG score indicates better axis alignment, meaning that a single latent variable is highly predictive of the factor, while others contribute minimally.

**TC** (Total Correlation)(Chen et al., 2018) assesses disentanglement in representation learning as a measure of statistical dependence among multiple random variables. It quantifies how much information is redundantly shared across latent dimensions, with a lower TC indicating greater independence.

The SAP score, MIG, and TC all evaluate disentanglement but from different perspectives. Together, they provide a comprehensive view of disentanglement—SAP and MIG focus on interpretability, while TC ensures minimal information overlap.

### 2.4 Expressiveness

Expressiveness is another quality of a representation and is evaluated by assessing its sensitivity to the removal of dimensions with redundant information, providing insights into how different dimensions contribute to model performance.

To measure expressiveness in representations, we incorporates two key concepts: (1) **Compactness** metric measures the ability of a small number of dimensions to effectively describe the representation manifold, aligning with the completeness aspect of DCI metrics. (2) **Intrinsic Dimension** captures the complexity of the representation manifold by estimating the number of parameters required to describe some downstream information without losing information (Thilak et al., 2023). This involves first training a model using the full set of dimensions, then removing a specified number of high-variance dimensions from the embeddings and evaluating performance on the reduced embeddings.

### 2.5 Robustness

Robustness to noise is a desired quality for latent representations, particularly in the context of generalization and stability. Robust latent representations should preserve meaningful structure while being insensitive to small perturbations or noise in the input data, making them more reliable for downstream tasks. The robustness test implemented in Latentverse evaluates various qualities, including clustering and downstream generalizability, across different levels of perturbation. This assessment helps determine how well the representation maintains its structure and performance under varying degrees of noise.

## 3. Latentverse

Our proposed library and web application called Latentverse, offers a suite of evaluation tests to assess the quality of latent representations from multiple perspectives.

### 3.1. Latentverse Code Library

Latentverse is released as an open-source library that provides a comprehensive suite of metrics, tests, and visualization tools for evaluating latent representations. It supports both standalone use and integration with the Latentverse web application. Users can install the package locally to run evaluations within their own workflows and easily incorporate it into their projects with minimal overhead. This enables seamless assessment of learned embeddings across a wide range of applications. The codebase is designed to be extensible, and we welcome community contributions, including the addition of new evaluation tests. To support this, the library includes a set of unit tests that help ensure correctness and reliability.

### 3.2. Latentverse Web Application Interface

Latentverse also offers an interactive web interface for evaluating learned representations. Users can upload representation files, along with optional task-specific labels, and select a test category from a drop-down menu. Latentverse then generates a comprehensive performance report that includes a suite of evaluation metrics corresponding to the selected test (Figure 1). For each metric, users can hover over the score to access detailed explanations—what the metric measures, how to interpret it, and its expected range.

**Figure 1.**
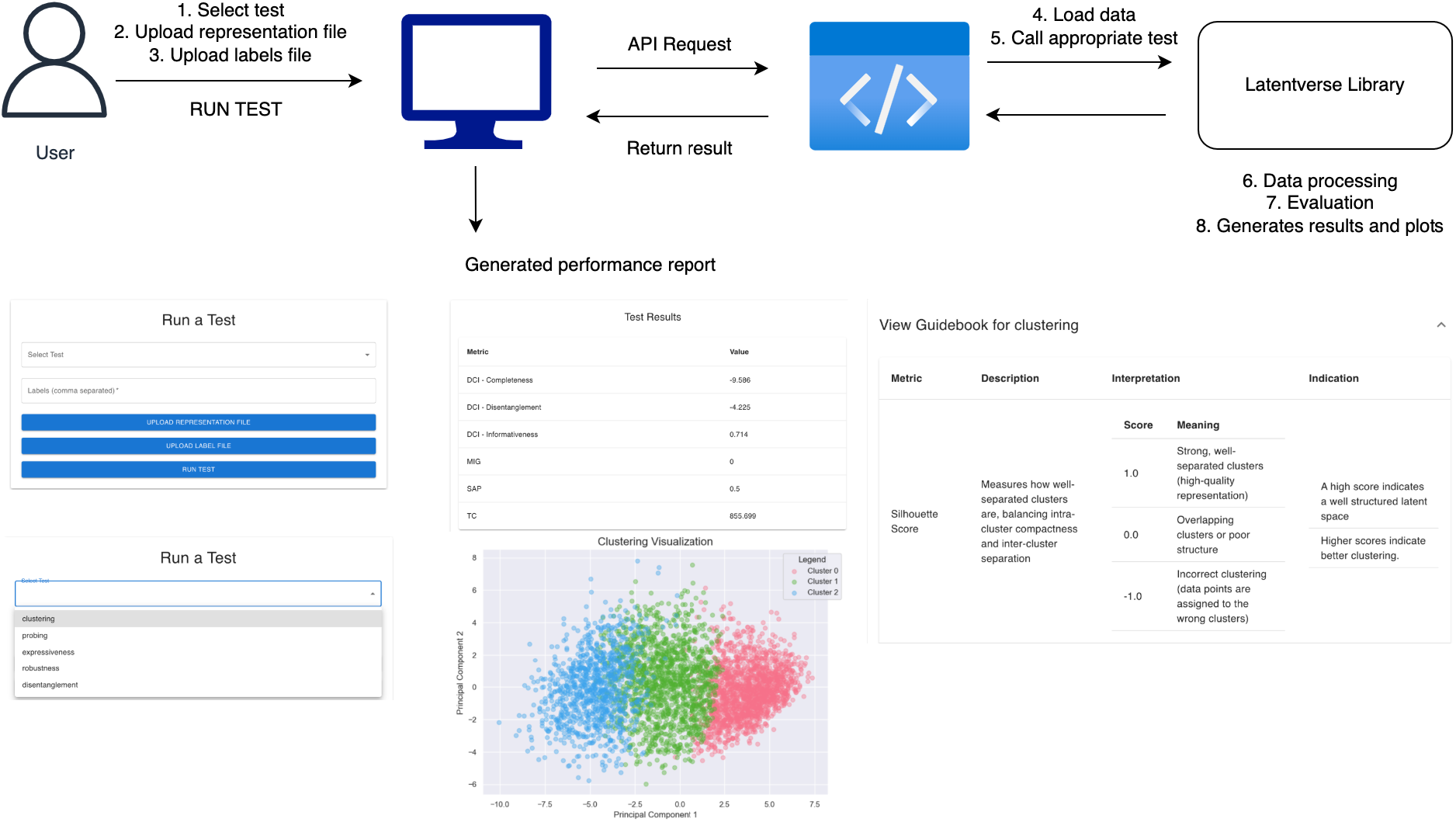
Overview of the Latentverse web interface. Users can upload latent representations for evaluation and select a test to generate a comprehensive report. The report includes various evaluation metrics and performance visualizations. For each test, there is a reference guidebook explaining the metric, interpreting the score values, and describing the implications for the representation. Additional information about each metric is available by hovering over it, providing details on what the metric measures, how to interpret it, and the corresponding range for evaluation.

To support interpretation, Latentverse includes a guidebook accessible via drop-down that provides structured insights for each metric. This guidebook explains the meaning of each score, its practical implications for representation quality, and offers recommendations for interpretation. As illustrated in Figure 1, the guidebook helps users contextualize results, enabling more informed decisions when refining or selecting embeddings.

In addition to metrics, Latentverse provides rich visualizations that highlight different properties of representations, aiding users in better understanding embedding quality. It also supports comparisons—allowing users to compare different representations or analyze the same representation under varying label sets for deeper insight.

### 3.3. Latentverse Web Application Implementation

The backend of Latentverse is powered by our opensource library and the web application is powered by a React-based frontend, a Flask-powered backend, and our open-source evaluation library. The application is deployed on a customizable Google Cloud Platform (GCP) virtual machine, allowing straightforward scaling based on user demand and uploaded file size. We can monitor resource usage over time and adjust the instance size accordingly, including adding a persistent disk for block storage if needed. Our setup uses Cloud Run, as its autoscaling capabilities could reduce maintenance overhead.

The frontend uses Material-UI for a design with interactive components such as the dropdown for test selection, representation and file upload fields, and hovering-activated pop-up text for describing each metric. Users are able to submit data, configure tests, and visual results using Matplotlib and Seaborn-generated plots that are displayed dynamically on the UI. Additionally, the Flask backend exposes a set of RESTful APIs that handle requests, data processing, and finally execute the evaluation tests. Upon user submission of data, the backend parses the request and performs validation on the input files, loads the representations and labels, calls the selected evaluation functions from the Latentverse library, and returns the computed performance metrics and corresponding visualization URLs to the frontend. Aside from the browser-based file upload we also support alternative solutions like Cloud storage integration, where users can first upload their files to GCP. Also, for the extreme cases where many users use the app at once, we have a load balancer in place to ensure the front end is stable.

## 4. Latentverse case-study

To demonstrate the utility of Latentverse, we conducted a study to show how Latentverse can be used to evaluate representations. We use a diverse set of latent representations learned from various data modalities and model architectures. Specifically, we use the UK Biobank dataset (Littlejohns et al., 2020), a large-scale biomedical resource containing detailed health and imaging data from over 500,000 participants in the United Kingdom. For our case study, we focused on electrocardiogram (ECG) and magnetic resonance imaging (MRI) modalities, training multiple encoder models to learn representations for these data types. We then evaluate and compare these representations using Latentverse to demonstrate how the tool can help users analyze the strengths and weaknesses of different models. Table 1 provides details about the latent representations used in our study, including information about training architectures and datasets, which serve as a basis for interpreting the evaluation results.

**Table 1:**
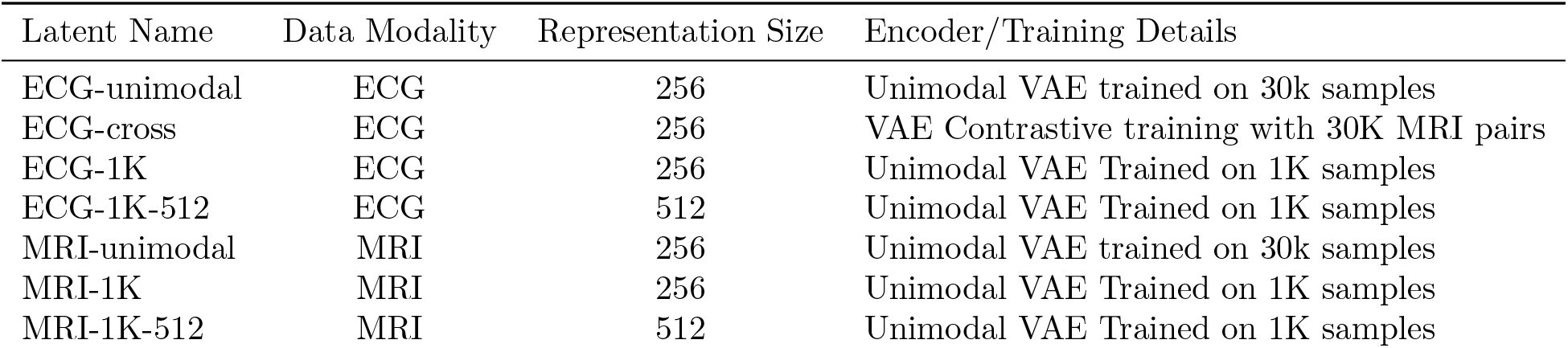
Details of the latent representations used for validating the Latentverse library. Data modality indicates the type of data used for learning embeddings. Representation size refers to the dimensionality of the embedding space. Encoder/training details describe the architecture of the self-supervised encoder and the training objective used to learn the representations.

| Latent Name | Data Modality | Representation Size | Encoder/Training Details |
| --- | --- | --- | --- |
| ECG-unimodal | ECG | 256 | Unimodal VAE trained on 30k samples |
| ECG-cross | ECG | 256 | VAE Contrastive training with 30K MRI pairs |
| ECG-1K | ECG | 256 | Unimodal VAE Trained on 1K samples |
| ECG-1K-512 | ECG | 512 | Unimodal VAE Trained on 1K samples |
| MRI-unimodal | MRI | 256 | Unimodal VAE trained on 30k samples |
| MRI-1K | MRI | 256 | Unimodal VAE Trained on 1K samples |
| MRI-1K-512 | MRI | 512 | Unimodal VAE Trained on 1K samples |

### 4.1. Downstream Generalization

To assess downstream generalizability, Latentverse measures probing performance for various labels using models with varying degree of complexity. We demonstrate this by comparing the predictive power of ECG-unimodal representations for estimating the RR interval against MRI-unimodal representations, which are learned from magnetic resonance imaging (MRI) data.

Figure 2 shows the results generated by Latentverse. RR interval is a key measure of the time between successive R-wave peaks in an electrocardiogram (ECG), which reflects heart rate variability. Therefore, it is a phenotype primarily captured by ECG data. We also see that the ECG-based representations exhibit higher probing performance compared to MRI-based representations for predicting this phenotype. We chose to display the R-squared metric here, but equivalently other measures can be selected. Interestingly, we observe variable probing performance depending on the complexity of the probing model. Simpler models are more interpretable and tend to generalize better, while more complex models achieve higher accuracy but can be prone to overfitting. This test provides insight into the amount of information a representation encodes about a certain label and how it can be used by downstream models.

**Figure 2.**
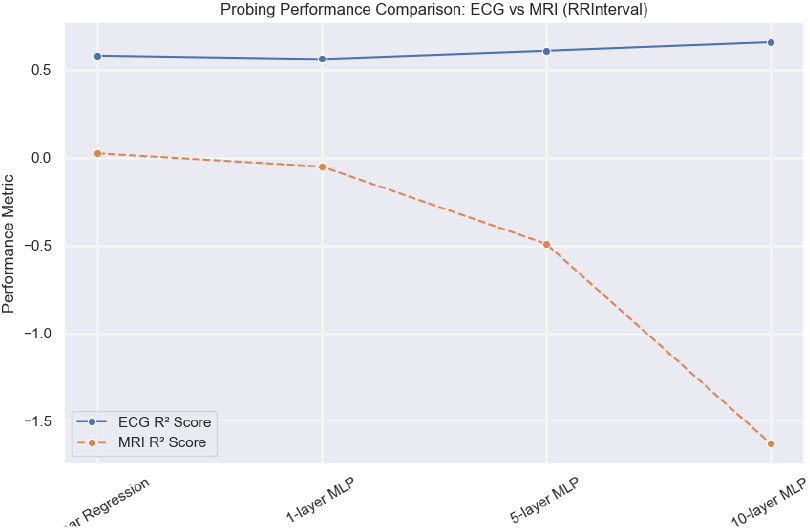
Downstream performance of ECG and MRI representations for estimating RR interval with probes of variable complexity.

### 4.2. Clusterability

To demonstrate the clusterability of representations, Latentverse provides a set of metrics along with visualizations that illustrate clustering behavior. Users have the option to include labels, enabling the tool to evaluate clustering performance based on those labels. Alternatively, Latentverse can assess the inherent clustering structure of the embedding space without relying on external labels.

Figure 3 shows the clustering performance report comparing ECG-unimodal and ECG-crossmodal representations for predicting patient biological sex. The cross-modal representation clearly captures sex-related information more effectively than the unimodal ECG embedding, suggesting that this information is primarily encoded through the MRI modality and learned theough the contrastive training. The NMI measure, which evaluates clusterability based on the specified labels, clearly shows this distinction. While some natural separation exists in the embedding space, there are no distinct clusters. As a result, metrics that rely on natural cluster separation, such as Silhouette and DBI, cannot distinguish the performance in this case. These scores are independent of evaluation labels, and simply measure the clustering behaviors.

**Figure 3.**
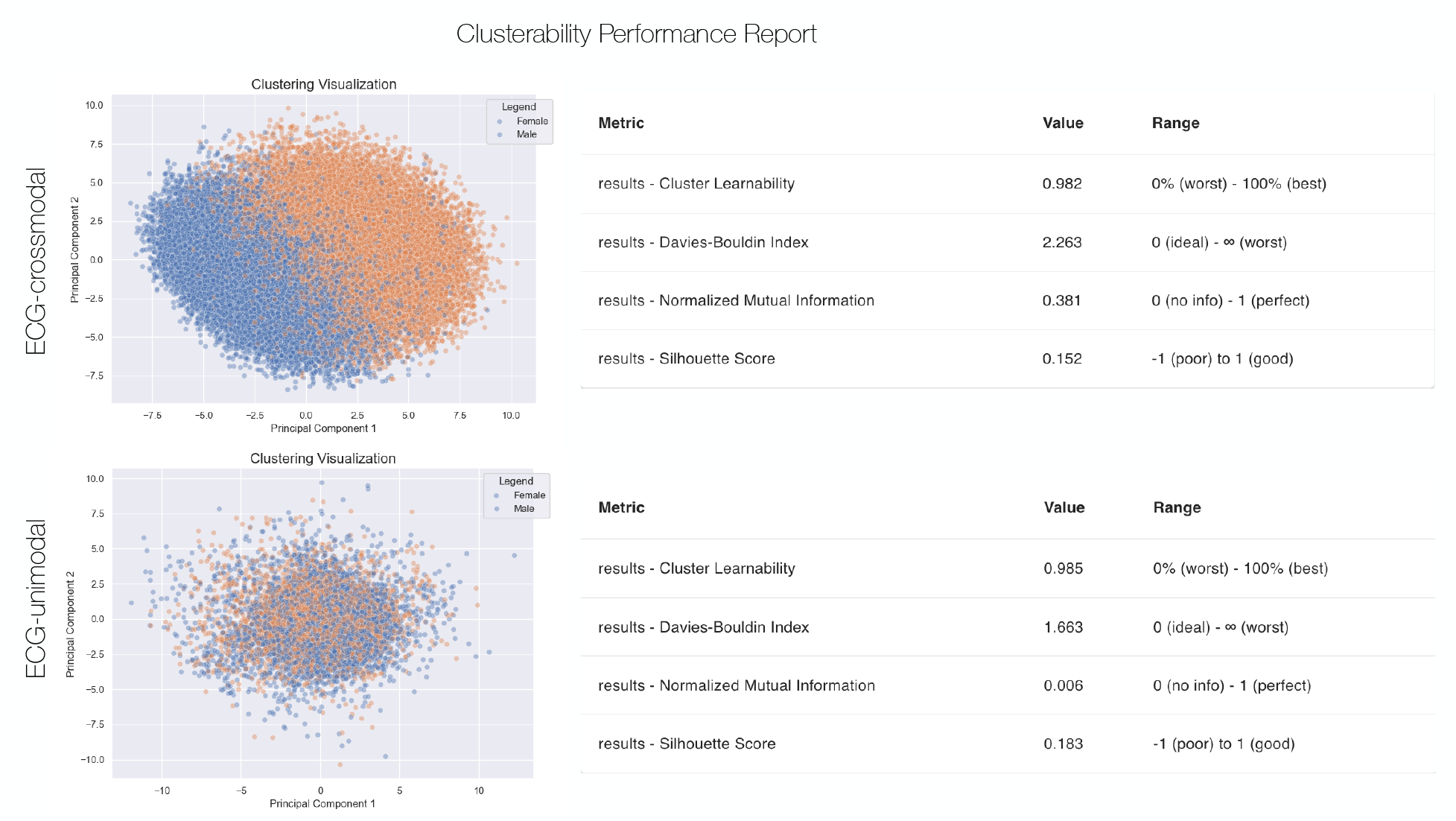
Clusterability performance report generated by Latentverse, which includes a number of evaluation metrics, as well as visual representations of the embedding space. This compares 2 sets of representations, ECG-unimodal and ECG-cross modal, comparing their clustering performance on separating samples based on sex. The information about MRI that is embedded in the cross-modal ECG representations enable a stronger separability.

### 4.3. Disentanglement

To demonstrate how Latentverse measures disentanglement, we modified one of our embedding spaces by adding an additional dimension that directly encodes a downstream label of interest—AFib (atrial fibrillation). Figure 4 shows the performance report for the ECG-unimodal representations and Figure 5 shows the performance for representations concatenated with AFib label.

**Figure 4.**
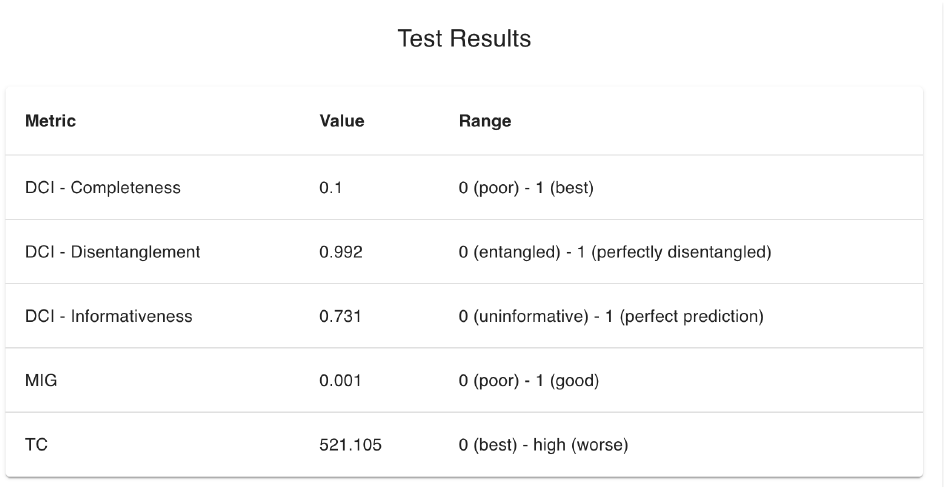
Disentanglement performance report on ECG-unimodal representations for estimating AFib.

**Figure 5.**
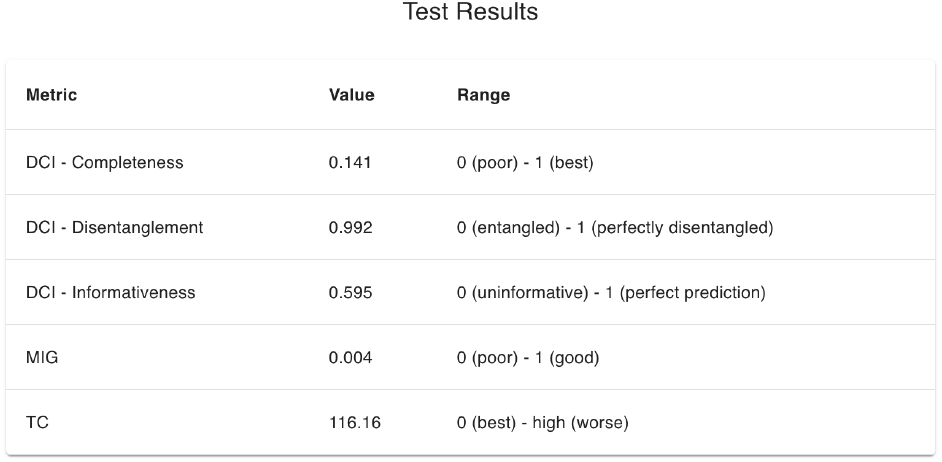
Disentanglement performance report on an augmented ECG-unimodal representations (with AFib labels) for estimating AFib.

As expected, adding the label information increases the MIG and completeness scores. The disentanglement score remains relatively unchanged, as it measures how distinct different generative factors are within the latent space, and other dimensions in the ECG space already capture information about AFib. Meanwhile, the high completeness score indicates that the AFib factor is now primarily represented by a single latent dimension. This highlights the importance of using a variety of metrics for assessing representations, as they provide complementary insights and enable a more thorough analysis of representation quality.

### 4.4. Expressiveness

Learning more compact representations is valuable, as larger embedding spaces are more prone to capturing noise and spurious correlations. The expressiveness test evaluates whether representations can be compressed into a lower-dimensional space with minimal loss of information. We applied this test on MRI representations with 512 dimensions (MRI-1K-512) and compared them to 256-dimensional representations (MRI-1K) for predicting biological sex. Figure 6 shows that for the higher-dimensional representations, removing up to 50% of the dimensions has minimal impact on performance, whereas the same level of removal significantly affects the lower-dimensional representations.

**Figure 6.**
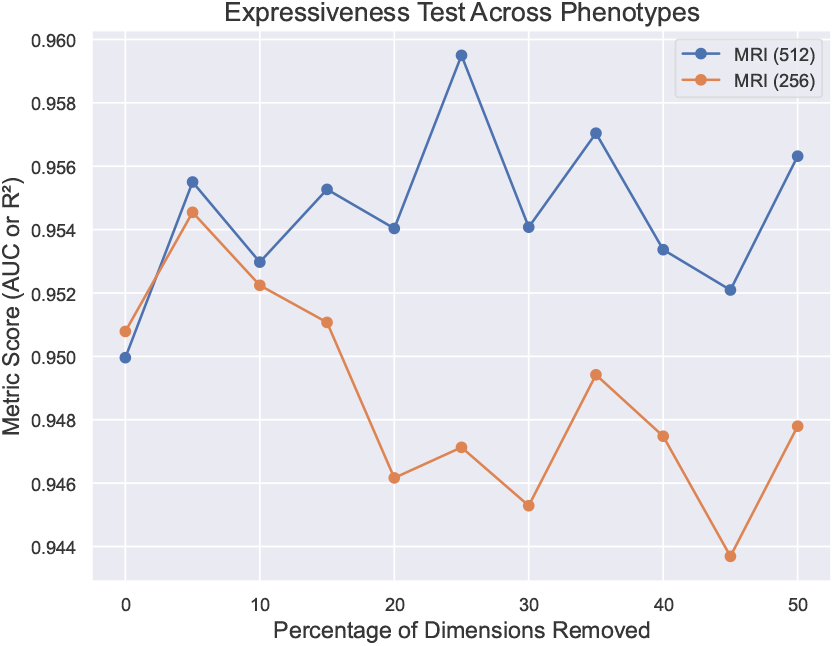
Expressiveness report comparing two sets of MRI representations with 256 and 512 dimensions. The plots show the predictive performance for predicting sex as a function of the percentage of highly correlated dimensions removed.

### 4.5. Robustness

Robustness tests typically focus on measuring sensitivity to perturbations in the input data. However, since our goal is to evaluate representations, we assess robustness to noise in the embedding space. This involves analyzing how the metrics discussed in previous sections respond to small perturbations in the representations. The test simulates data corruption by introducing Gaussian noise into the latent space and observing how performance metrics are affected, providing insights into the stability and resilience of the learned representations. Figure 7 shows an example of the performance report Latentverse generates for the ECG-unimodal representation. Overall, with increasing noise, the clustering performance worsens. We see how the DBI is more sensitive to noise compared to other metrics due to the impact of noise to increased intra-cluster variance and reduced intercluster distance. Silhouette Score focuses on individual point separation, making it more robust to local noise.

**Figure 7.**
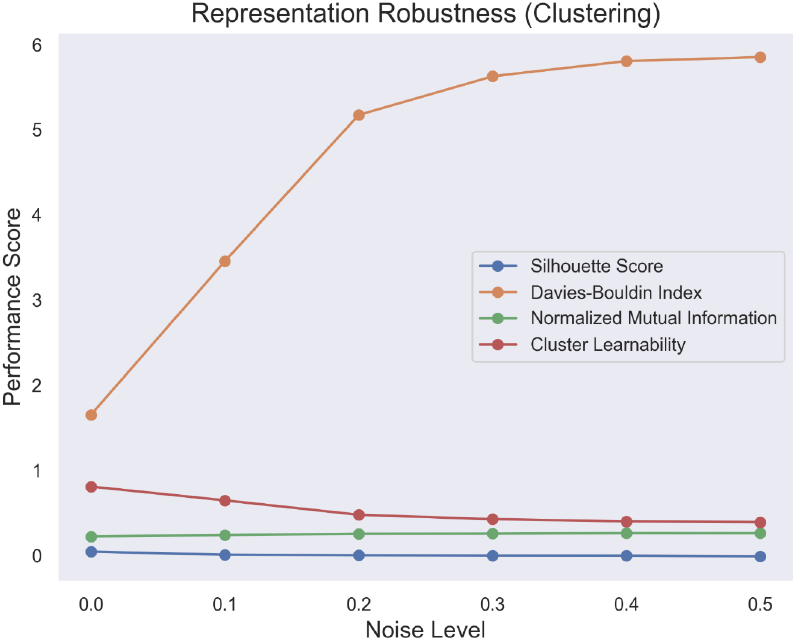
Robustness of clusterability performance on ECG-unimodal representations as a function of varying noise levels.

Figure 8 shows a different example using the ECG1K model (an ECG encoder trained on 1K samples), comparing a 256-dimensional representation against a 512-dimensional one. Unless carefully regularized, higher-dimensional embeddings can include redundant or uninformative dimensions that increase sensitivity to noise. While the initial performance of the 256-dimensional representation is lower than that of its higher-dimensional counterpart, it demonstrates greater robustness under increasing noise. In fact, under more extreme noise conditions, the 256-dimensional representation eventually outperforms the 512-dimensional one, likely due to its lower capacity and more concentrated feature encoding.

**Figure 8.**
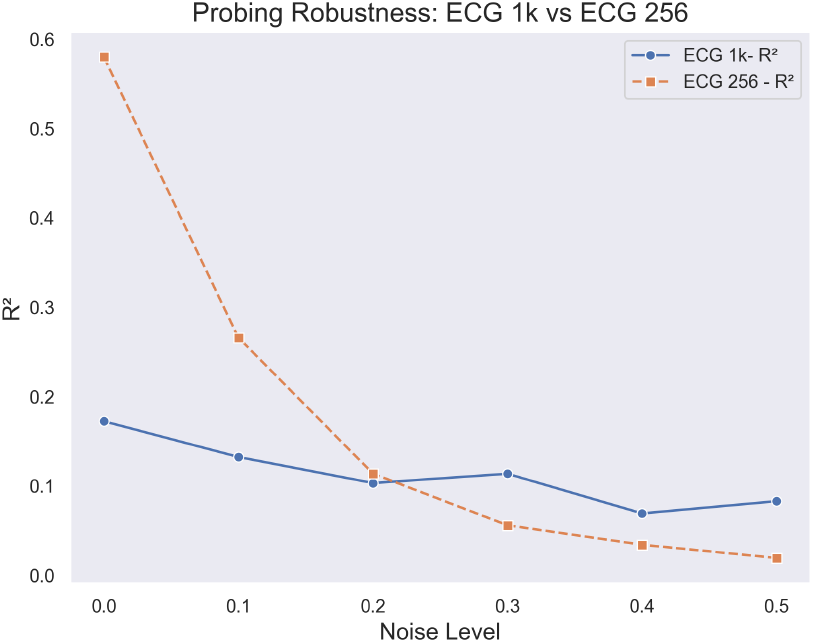
Robustness of probing performance on ECG representations with 512 and 256 dimensions as a function of noise levels

**Figure 9.**
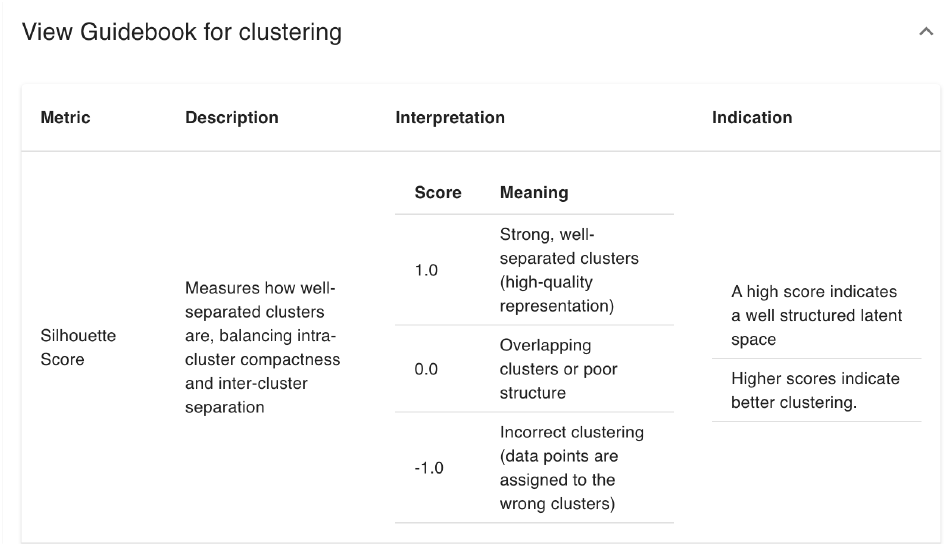
Guidebook for clustering test results.

**Figure 10.**
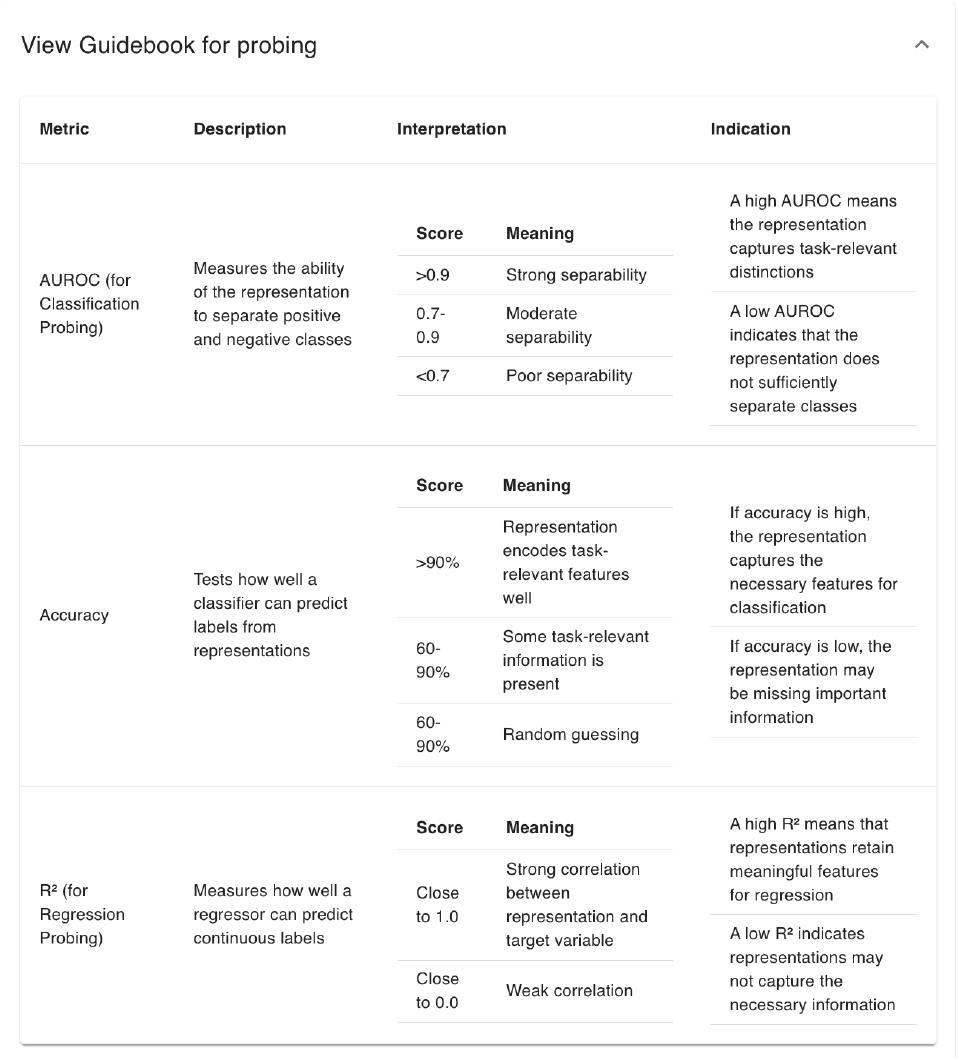
Guidebook for probing test results.

**Figure 11.**
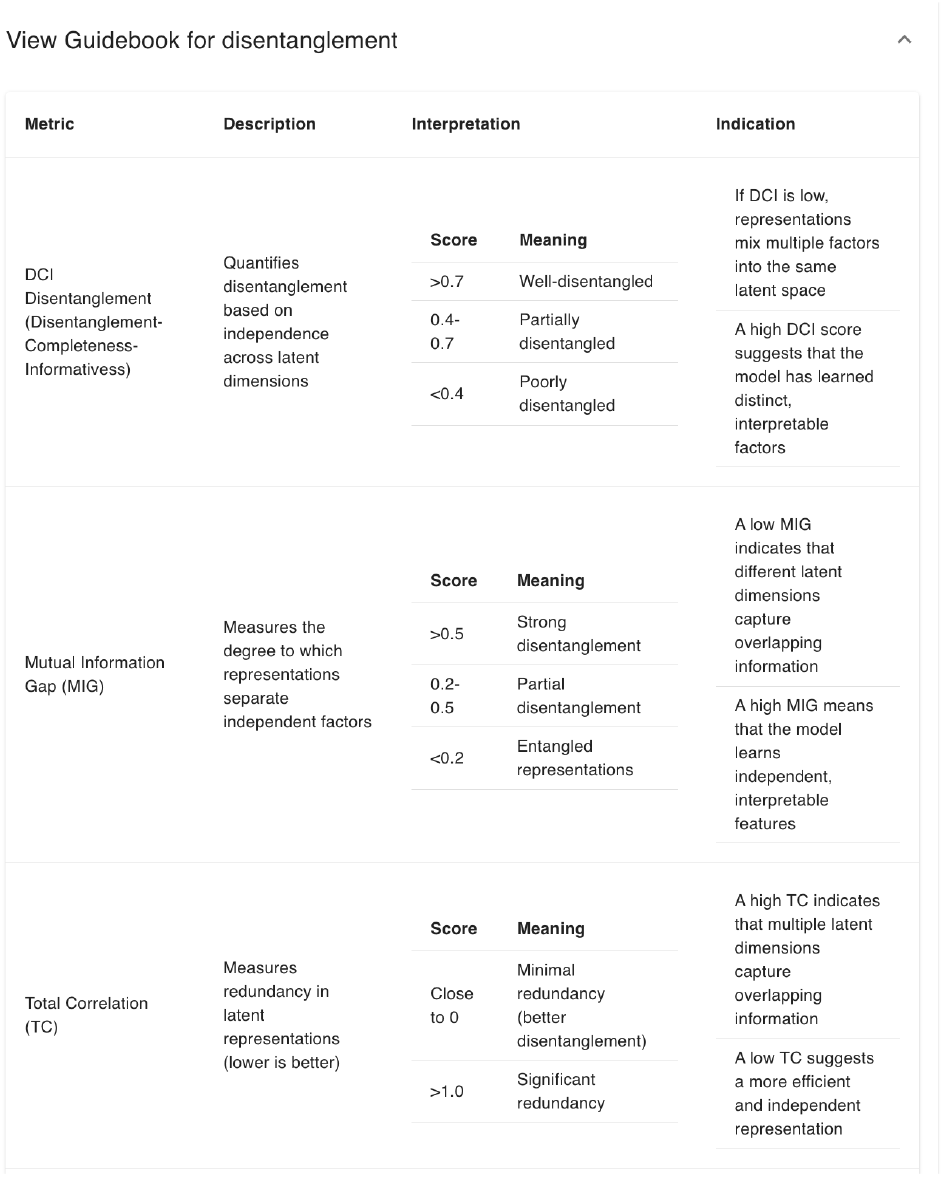
Guidebook for disentanglement test results.

## 5. Discussion

With the growing adoption of self-supervised learning and pretrained models, evaluating the quality and behavior of latent representations has become increasingly crucial. In this paper, we introduce Latentverse, a tool that provides a multi-faceted evaluation framework to give users a comprehensive understanding of representation quality. Latentverse enables users to analyze and interpret representations, assess how they perform across different scenarios, and make more informed model selection and finetuning decisions. By offering both evaluation metrics and comparison tools, Latentverse supports deeper insights into latent representations.

Future work will focus on expanding the interface to offer more automated result analysis, making the tool accessible to users from diverse technical backgrounds. We share Latentverse as an open-source library, aiming for it to become an evolving resource that meets the needs of this fast-growing field. We hope Latentverse becomes a collaborative platform where researchers and practitioners can contribute new evaluation metrics and develop innovative ways to assess representation quality.

## Acknowledgments

This work has been supported by the Eric and Wendy Schmidt Center at the Broad Institute of MIT and Harvard. The authors thank the UK Biobank for providing the data for this study. Data were accessed under the Broad Institute’s UK Biobank application number 7089

## Appendix A. Web Application Interface

### A.1. Guidebook

Latentverse evaluates representations using a wide range of metrics. Each metric measures different aspects of the representations and will have its own strengths and weaknesses. In order to help users in their analysis of the results, Latentverse provides a guidebook page that explains the nuances of different metrics, what different values indicate, and their strength and weaknesses for different scenarios.

